# Decisions in an Innate Behavioral Sequence

**DOI:** 10.1101/2021.04.03.438315

**Authors:** Kevin M. Cury, Richard Axel

## Abstract

Innate behaviors are comprised of ordered sequences of component actions that progress to satisfy drives. We have characterized the structure of egg-laying behavior in *Drosophila* in detail and observed that the sequence is not merely comprised of motor acts but also acts of sensory exploration that govern the transitions between component actions. We have identified a cluster of internal sensory neurons that provide information about the progression of the egg during ovipositor burrowing, a behavior necessary for the subterraneous deposition of the egg. These neurons impart sensory feedback that allows burrowing to continue to egg deposition or to abort in favor of further exploration. Diminished activity of these neurons upon completed egg expulsion may initiate the transition to the final phase of egg-laying, allowing the cycle to repeat. Sensory feedback therefore plays a critical role at decision points between transitions affording innate behaviors with an adaptive flexibility.

## INTRODUCTION

Organisms have evolved a repertoire of innate behaviors, comprised of sequences of component actions, to satisfy essential drives ^1–3^. Innate behaviors are often considered fixed or stereotyped, but the individual component actions can exhibit considerable variation ^4–6^. Variability in an innate behavior is greatest during an initial appetitive phase during which the organism is seeking an appropriate environment or social context ^7^. This is followed by a consummatory phase comprised of conserved actions that directly satisfy the drive. Behavioral variability results from the acquisition of sensory feedback that informs decisions at the junctures between component actions. The presence of such decision points may allow an assessment of progress and afford the organism with adaptive flexibility to accommodate variability in the internal and external environment.

The drive to reproduce is a dominant motivator of behavior in all species. Diverse behavioral programs dedicated to courtship, copulation and the production and care of offspring have evolved to optimize reproductive success. For oviparous animals that do not brood, such as the fruit fly, *Drosophila melanogaster*, egg deposition represents the culmination of this array of reproductive behaviors. Considerable pressure is imposed on the selection of the appropriate time and place to deposit eggs. Fruit flies express robust and species-specific preferences for the site of egg deposition based in part on odor, taste, and texture ^8–15^. During egg-laying, females evaluate the local environment prior to expressing an ordered motor sequence that culminates in egg deposition subterraneously within a nutritive substrate ^14,16–18^. Upon egg deposition the female comes to rest and this final phase of the behavioral sequence is coupled to ovulation and fertilization, and the cycle repeats.

Innate behavioral programs are orchestrated by genetically determined neural circuits. Higher order centers in the brain integrate internal and external stimuli that reinforce a representation of a dominant drive^1,16,19–24^. These centers activate lower level circuits in the ventral nerve cord or spinal cord that elicit organized sequences of behavior ^25–28^. These lower level circuits intrinsically generate coordinated motor patterns but sensory feedback is critical in adapting these patterns to accommodate variability in the environment.

The neural circuit responsible for the sequence of egg laying behaviors in *D. melanogaster* is activated following mating. The drive to lay eggs is elicited by a seminal fluid peptide, sex peptide, introduced into the uterus during mating ^29^. Sensory neurons responsive to sex peptide transmit this information to the brain, and inhibit a subset of the pC1 cluster of neurons ^16,30–32^. This disinhibits oviDN, a cluster of descending interneurons that project to the ventral nerve cord and are necessary and causal for the expression of the egg deposition motor sequence. OviDN may be a critical intermediate between higher order centers that represent egg-laying drive and the motor circuits that execute the terminal sequence of egg-laying behavior. However, progression along this terminal motor sequence and the successful deposition of the egg may be dependent on sensory feedback.

We have characterized the detailed structure of egg-laying behavior and observe that it consists of variable transitions between component actions prior to egg deposition. Our analysis demonstrates that the behavioral sequence is not merely comprised of motor acts but also reveals acts of sensory exploration that shape the motor program. We have identified a cluster of internal sensory neurons that provide feedback regarding the progression of the egg during ovipositor burrowing, the concluding act necessary for the subterraneous deposition of the egg. The activity of these neurons provides feedback that allows burrowing to continue to egg deposition or to abort in favor of further exploration. Diminished activity of these neurons upon completed egg expulsion results in the transition to the final phase allowing the cycle to repeat. Flexibility is therefore imposed on an innate and stereotyped sequence of motor actions to optimize the deposition and successful development of the egg.

## RESULTS

### Egg-laying sequence exhibits variable transitions between component actions prior to egg deposition

Females lay eggs one at a time in a repeating cycle, continually transitioning between three distinct phases of a behavioral sequence. Each egg-laying cycle is comprised of an active exploratory phase, deposition, and a more stationary phase (“reset”) that includes ovulation, after which the cycle repeats ^14,18^ (Figures 1, S1A and S1B). We have studied the component behaviors of this sequence in detail by filming individual gravid females at high resolution in small egg-laying chambers (Figure S1C and Supplemental Video S1). Prior to deposition, flies explored the substrate with their legs and proboscis. During this phase, flies walked, paused, and extended their proboscis to make brief contact with the substrate (“PE”, proboscis extension, in Figure 1A). They then bent their abdomen to bring the ovipositor in contact with the surface (“bend” in Figure 1A). The flies then transitioned to deposition, and initiated substrate burrowing – a rhythmic behavior in which the ovipositor digs into the substrate and expels the egg – and ultimately deposited the egg subterraneously (“burrow” and “egg out” in Figure 1A). Upon completed egg deposition, the flies abruptly stopped burrowing, detached from the egg and then lifted and groomed their ovipositor (“detach” and “groom” in Figure 1A). The females then remained stationary for an extended period of time, intermittently grooming and exhibiting abdominal contortions likely to result from ovulation (the “reset” phase). The behavioral sequence then repeated. This ordering of component actions was highly conserved across repeated egg-laying events (Figure 1B). Thus, egg-laying behavior appears to be organized as an ordered sequence of behavioral components.

**Figure 1.**
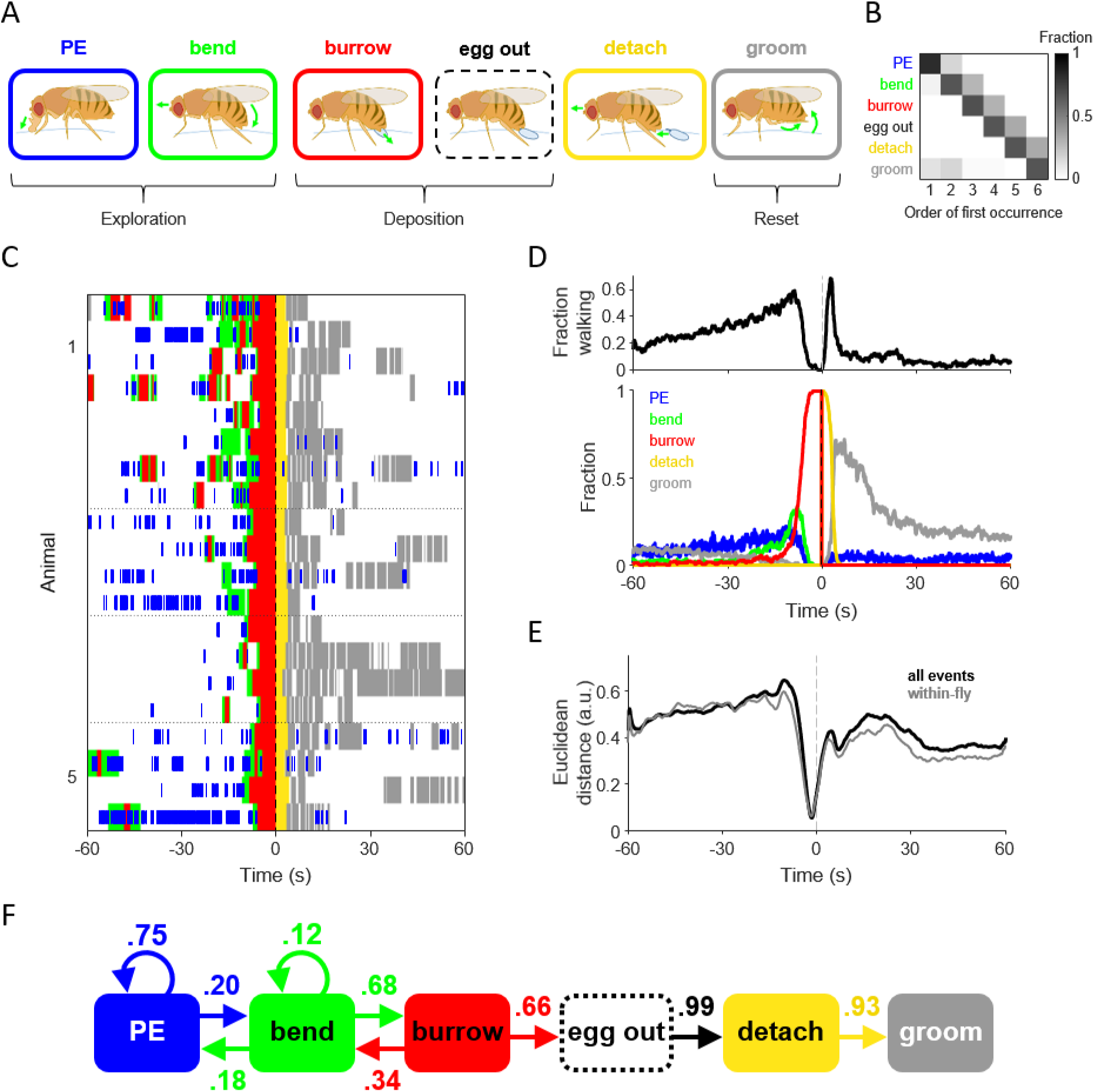
Egg-laying sequence exhibits variable transitions between component actions prior to egg deposition. (A) Illustrations depicting component actions of the egg-laying behavioral sequence. Components comprising exploration, deposition, and “reset” phases are indicated. The colors used in text and boxes are also used to indicate component actions in all subsequent figures. “PE”, proboscis extension. (B) The order of occurrence of the first instance of each behavioral component, depicted as a fraction of total events. Only events including all components were analyzed (169 of 176 events). (C) Ethogram of egg-laying behavior in five flies (n = 4 events per fly). surrounding egg deposition, color-coded as indicated in (A). Time = 0 marks the time of completed egg deposition (“egg out”) here and in (D) and (E). (D) Top, fraction of flies in motion at a given time surrounding egg deposition. Bottom, time course of the five annotated behaviors. n = 176 egg-deposition events from 18 flies. (E) Black trace, average Euclidean distance between all pairwise combinations of behavioral trajectories, expressed as a 6-dimensional vector (the five annotated behaviors and movement). Gray trace, average Euclidean distance between events in the same fly. Behavioral trajectories were smoothed with a 5-second moving average. (F) Diagram depicting the transition probabilities between the five annotated behaviors and egg deposition (“egg out”).

However, we observed considerable variability in the timing, frequency, and duration of the individual components within this behavioral sequence. Variability was most apparent during the exploration phase. In contrast, the behaviors surrounding egg deposition formed a sequence (burrow - egg out - detach - groom) that was invariant (Figures 1C and 1D). We determined the Euclidean distance between behavioral trajectories over a sliding window and observe maximal separation (behavioral divergence) during exploration (Figure 1E). During this phase, flies were most likely to perform different actions at a given time. Intermediate separation was observed during the “reset” phase. In contrast there was striking convergence of behavior surrounding egg deposition. These trends were observed across repeated egg-laying events in the same fly and across events in the population of flies (Figure 1E, gray trace versus black trace).

We more closely examined the ordering of component actions by determining the probabilities of transitions between components along the egg-laying sequence (Figure 1F). During exploration and deposition, the behavioral sequence was conserved but transitions could occur in both directions. Although burrowing was always preceded by abdominal bending, bending was not always followed by burrowing. Rather, additional bouts of bending and even proboscis contact were observed. Thus, individual component behaviors can be repeatedly expressed in new locations or the fly can revert to a preceding behavioral component. This variability in transitions was not only apparent for proboscis and ovipositor contact (“PE” and “bend”), but was also observed for burrowing. For example, 34% of burrowing episodes did not persist to egg deposition but were aborted in favor of further exploration. Persistent burrowing invariably preceded egg deposition and, in accord with our measurement of Euclidean distance, behavioral transitions following egg deposition exhibited little variability and proceeded along the ordered sequence with high probabilities to the “reset” phase. Thus, prior to the completion of egg deposition, the egg-laying sequence is comprised of multiple junctures between component actions that may serve as “decision points.” These junctures may allow the fly to advance or suspend the sequence contingent on sensory information obtained during a component action.

### Burrowing behavior is flexibly adjusted to changes in substrate firmness

Burrowing precedes the final “decision point” in the egg-laying sequence: egg deposition and the transition to the “reset” phase. This pivotal role led us to closely examine the sub-structure of burrowing. A burrowing episode is comprised of discrete cycles that begins with rhythmic ovipositor digging (Figure 2A). As the surface is scored, the ovipositor extends into the substrate and the egg emerges out of the uterus and into the ovipositor. Rhythmic pushing expels the egg out of the ovipositor and into the substrate, just beneath the surface. Completed egg deposition halts the rhythm, terminating the burrowing episode, and the fly then detaches the ovipositor from the egg.

**Figure 2.**
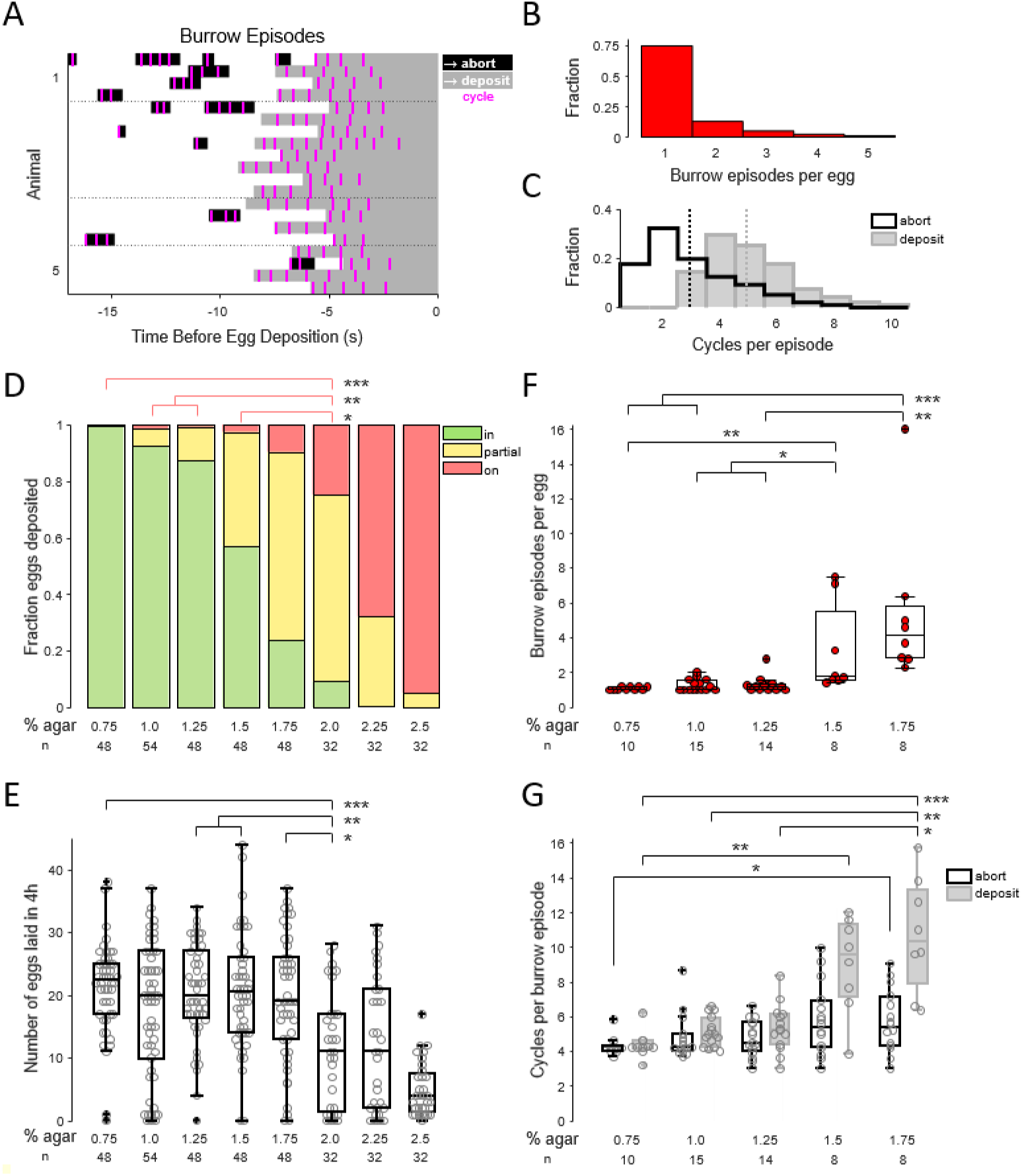
Burrowing behavior is flexibly adjusted to changes in substrate firmness. (A) Ethogram depicting burrowing episodes in five flies (n = 4 events per fly). Black, aborted episode; gray, egg-deposition episode; magenta, cycles within a burrowing episode. Time = 0 marks the time of completed egg deposition (“egg out”). Note that these data are the same events depicted in Figure 1C. (B) Distribution of the number of burrowing episodes per egg-deposition event. Data here and in (C) are from experiments described in Figure 1 and are pooled across all flies. (C) Distributions of the number of cycles per burrowing episode. Black, aborted episode; gray, egg-deposition episode. The mean value for each distribution is indicated by a dashed vertical line. (D) Stacked distributions of the average depth of penetration of eggs laid across substrates of increasing firmness (see Methods). The mean fraction within a group is presented here. *p<.05, **p<0.01, ***p<0.001, Kruskal-Wallis test with post hoc Tukey’s HSD test here and in E-G. Significance only depicted for comparisons between fraction deposited “on” the substrate for groups from 0.75% to 2.0% agar. (E) Number of eggs laid across substrates of increasing firmness in a 4-hour window. In all subsequent plots, box represents the 25^th^ and 75^th^ percentile with the median indicated by a horizontal line, and whiskers represent the 5^th^ and 95^th^ percentile. Outliers are indicated with a and individual data points with a “o.” Significance only depicted for comparisons up to 2.0% agar. (F) Average number of burrowing episodes per egg-deposition event across substrates of increasing firmness. Only episodes occurring on the substrate and not the chamber walls were considered here. (G) Average number of cycles per burrowing episode for both aborted and egg-deposition episodes across substrates of increasing firmness.

We observed, however, that not all burrowing episodes persisted to egg deposition. Episodes could be aborted and re-initiated at another location. In chambers containing 1% agar, 75% of the eggs were deposited during the first burrowing episode, whereas the remaining eggs were deposited following at least one aborted episode (Figures 2A and 2B). 73% of aborted episodes occurred when the fly was on the rigid chamber walls and 97% of episodes on the wall were aborted. Egg deposition required a minimum of 3 cycles and could require as many as 10 cycles within an episode (Figure 2C, gray histogram). The number of burrowing cycles was significantly lower in aborted episodes (mean of 3 cycles for aborted, 5 cycles for deposition; p<0.001, Wilcoxon rank sum test), and burrowing could be aborted after any cycle within an episode (min of 1 - max of 8; Figure 2C, black histogram). This suggest that the decision to persist in burrowing may be determined after each individual cycle. Thus, the number of burrowing episodes and the number of cycles within an episode necessary to deposit an egg showed considerable variability. Burrowing can therefore be repeated or extended to achieve successful egg deposition.

The variability in the number of burrowing episodes and the number of cycles within an episode may reflect variability in the properties of the egg-laying site. We therefore asked whether burrowing adapts to changes in the firmness of the substrate ^9,10,33^. We scored both the total count and the depth of penetration of eggs laid on agar substrates of increasing firmness (agar concentrations from 0.75% to 2.5%; Figures 2D and 2E). As substrate firmness was increased, flies were less successful at achieving subterraneous egg deposition (Figure 2D). Above 1.75% agar, total egg output dropped significantly (mean of 11 eggs laid in 4h on 2.0%, compared to 18-21 eggs for 0.75% to 1.75%; Figure 2E) and flies deposited a significantly larger fraction of eggs on the substrate surface (Figure 2D). This suggests that the ability to achieve subterraneous egg placement positively gates egg output.

The subterraneous deposition of an egg is sensitive to substrate firmness and depends upon burrowing, suggesting that burrowing behavior will be modified in response to change in substrate composition. We observed that the structure of burrowing behavior was dramatically altered as the firmness of the agar substrate was increased. The number of burrowing episodes per egg increased five-fold and the total number of burrow cycles required for egg deposition increased over two-fold (Figures 2F and 2G) as the agar concentration increased from 0.75% to 1.75%. Thus, additional burrowing cycles are required to dig and push the egg into the firmer substrates. If the egg cannot be successfully deposited, burrowing is aborted in search of a more favorable location. These data suggest that the transition to egg deposition and the “reset” phase is contingent on the decision to persist in burrowing until the egg is completely expelled. The decision to persist in burrowing is likely to be informed by ongoing sensory feedback regarding the progression of the egg as it is pushed through the uterus and ovipositor into the substrate.

### PU sensory neurons are activated by the presence of the egg in the ovipositor

We next screened a library of LexA lines ^34^ to identify candidate sensory neurons that detect the progression of the egg ^12,35–37^. We identified three lines that drive expression in a cluster of peripheral neurons whose processes intercalate with muscle fibers that encircle the distal-most aspect of the uterus (Posterior Uterine sensory, or PU neurons) (Figure S2A). We used the split-GAL4 intersectional strategy using enhancers from these three LexA lines to produce lines with restricted expression in PU neurons ^38,39^. Two of these split-GAL4 lines (hereafter referred to as PU-1 and PU-2) typically labelled a pair of PU neurons on each side of the uterus that extended processes into the ventral abdominal ganglion (PU-1, 1.9 ± 0.5 cells per side, n = 17; PU-2, 2.1 ± 0.3 cells, n = 11; mean ± SD; Figures 3A, 3B, and S2D-F). No labelling was observed in the brain though, in addition to PU neurons, both lines exhibited inconsistent labelling in a small number of peripheral neurons that project to the dorsal abdominal ganglion (PU-1, 1.2 ± 0.6 dorsal afferents per side; PU-2, 0.5 ± 0.6; mean ± SD). We confirmed that PU-neurons are indeed sensory neurons by targeted expression of the somatodendritic marker DenMark which localized to the peripheral processes encircling the posterior uterus (Figure S2E) ^40^. This pattern of dendritic innervation suggests that PU neurons may sense the passage of an egg from the uterus to the ovipositor.

**Figure 3.**
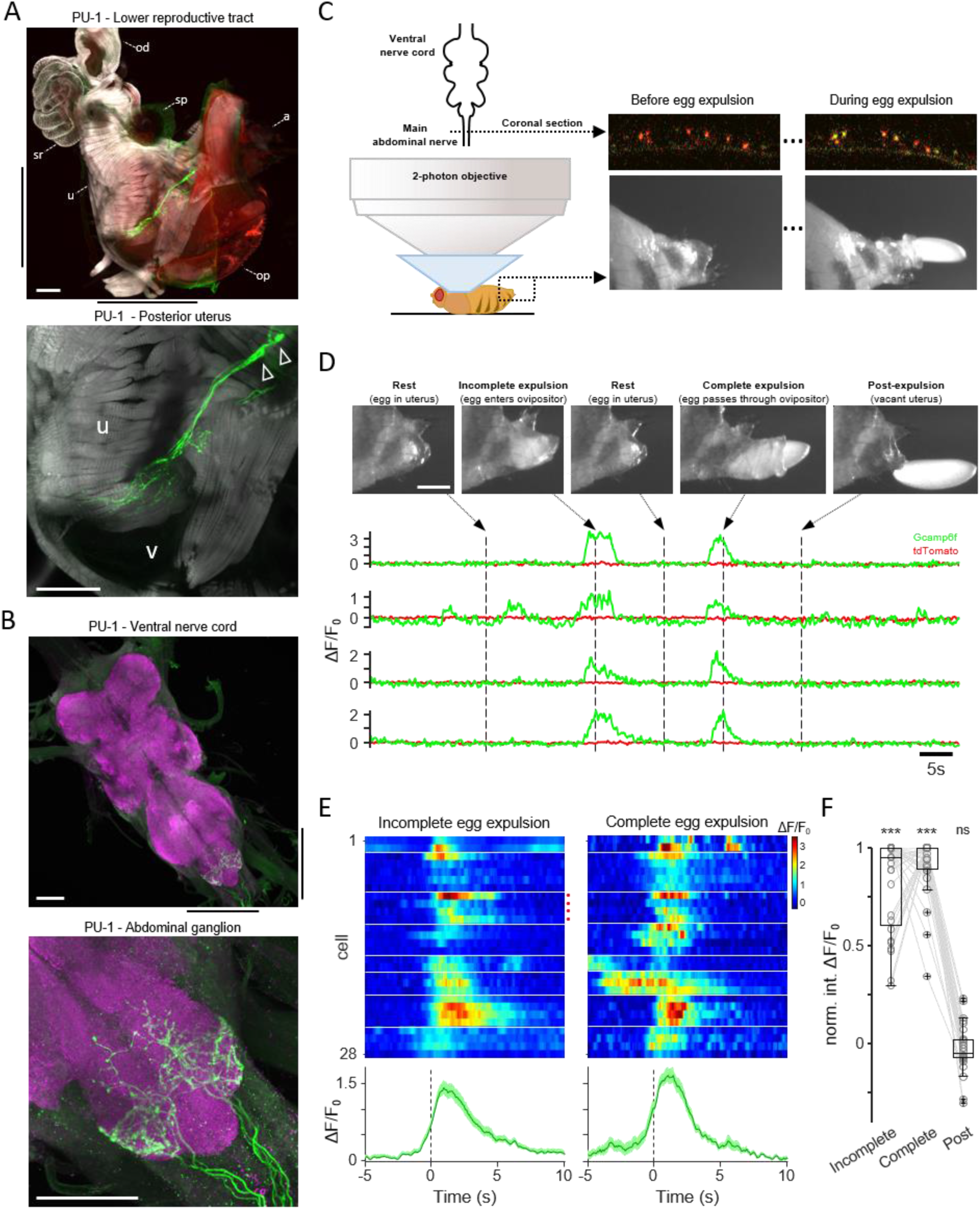
PU sensory neurons are activated by the presence of the egg in the ovipositor. (A) Top, confocal image of the lower reproductive tract and posterior abdomen from a PU-1 > CD8-GFP female, stained with anti-GFP to reveal the membrane of PU neurons (green) and anti-F-actin to visualize muscle fibers (gray). The abdominal cuticle is visualized by autofluorescence (red). Black bars outside the top image indicate the region shown in higher resolution in the bottom image. ‘od’, oviduct; ‘sp’, spermathecae; ‘a’, anal plates; ‘sr’, seminal receptacle; ‘u’, uterus; ‘op’, ovipositor. Bottom, two PU cell bodies (white triangles) and their dendrites. ‘u’, uterus; ‘v’, vagina. Scale bar is 50 μm. (B) Confocal images of the ventral nerve cord (top) and abdominal ganglion (bottom) from a PU-1 > CD8-GFP female, stained with anti-GFP (green) and nc82 (magenta) to visualize the entire synaptic neuropil. Black bars outside the top image indicate the region shown in higher resolution in the bottom image. Scale bar is 50 μm. (C) Two-photon experimental setup allowing for the simultaneous measurement of Gcamp6f (green) and tdTomato (red) fluorescence in axons within a coronal section of the main abdominal nerve (top right panels) and videography of the posterior abdomen (bottom right panels). (D) Top, video snapshots of the posterior abdomen at multiple time points including incomplete and complete egg expulsion, recorded concurrently with two-photon imaging. Scale bar is 200 μm. Bottom, two-photon imaging of four PU axons, encompassing the above events, depicting relative fluorescence changes (ΔF/F_0_). Green traces, GCamp6f fluorescence; red traces, tdTomato anatomical marker. Arrows and vertical dashed lines indicate the corresponding time point for each video snapshot presented above. Note that the video snapshots were flipped in the vertical dimension here. (E) Relative fluorescence changes in PU neurons in response to incomplete egg expulsion events (left) and complete egg expulsion (right), with individual responses shown on top and the average response across all neurons shown on bottom. Bottom, darker traces indicate mean response; lighter area represents SEM. The cells from (D) are indicated by a red dot. If multiple incomplete egg expulsion events occurred, the average response per neuron is presented here (n = 54, 1, 2, 5, 1, 1, 4, 5 events per fly). Horizontal white lines demarcate recordings performed in different flies. Time = 0 is the time of event onset (see Methods). (F) Normalized population data, showing the 3-second integrated ΔF/F_0_ fluorescence levels during incomplete egg expulsion, complete egg expulsion, and post egg-expulsion. Statistical comparisons were made with the pre-expulsion baseline, “0” value. ***p<0.001, ns p>0.05, Wilcoxon signed rank test.

We therefore developed a fly preparation that allowed us to monitor the activity of PU neurons as the egg is expelled from the uterus (Figure 3C and Supplemental Video S2). GCamp6f was expressed in PU neurons, and calcium activity was recorded by two-photon imaging of PU axons within the main abdominal nerve ^41,42^. Infrared video recordings of the exposed posterior abdomen allowed synchronous visualization of the presence and position of the egg. Snapshots from a typical video recording over a 90-second window encompassing egg expulsion are shown in Figure 3D, along with the corresponding activity of the PU axons. Initially, we observe an incomplete expulsion event (Figure 3D, 2nd frame): the egg progresses from the uterus (Figure 3D, 1st frame) into the extruded ovipositor (Figure 3D, 2nd frame), after which the egg retreats into the uterus and the ovipositor retracts (Figure 3D, 3rd frame). During this event, calcium activity in PU neurons increases from baseline when the egg enters the ovipositor and returns to baseline when the egg retreats into the uterus. A second, complete expulsion event then occurs, and the PU neurons again respond when the egg enters the ovipositor, and the activity returns to baseline after the egg is fully expelled (Figure 3D, 4th frame and 5th frame). These response properties were observed in all 28 PU neurons recorded from 8 flies (Figures 3E and 3F). Interestingly, PU neurons were not activated by the mere presence of the egg in the uterus (Figures 3D-F) nor did they respond to ovipositor extrusion events in flies lacking an egg (Figures S2F-H). These observations demonstrate that PU neurons respond when the egg progresses from the uterus into the ovipositor. Thus, PU neurons are poised to inform circuits in the abdominal ganglion about the progression of the egg during burrowing.

### Silencing PU neurons reduces egg output and disrupts multiple components of the egg-laying sequence

We next chronically silenced PU sensory neurons to examine their role in egg-laying behavior by targeted expression of the potassium channel, Kir2.1 ^43^. In PU-silenced flies we observed a dramatic reduction in the number of eggs laid (PU-1 >Kir2.1, mean of 9 eggs laid in 4h compared to 23 and 34 in genetic controls; Figures 4A and S3A). Moreover, the majority of these eggs were not deposited subterraneously on a 1.0% agarose substrate (PU-1>Kir2.1, 29% deposited subterraneously versus 95% and 93% in genetic controls; Figures 4B and S3B). This reduction in egg count was not a consequence of a mating defect. PU-silenced virgins showed no deficit in mating, but exhibited a reduction in egg count after a single mating event (Figures S3C and S3D). Interestingly, half of the eggs laid did not hatch suggesting that PU-silencing also compromises fertilization (Figure S3E).

**Figure 4.**
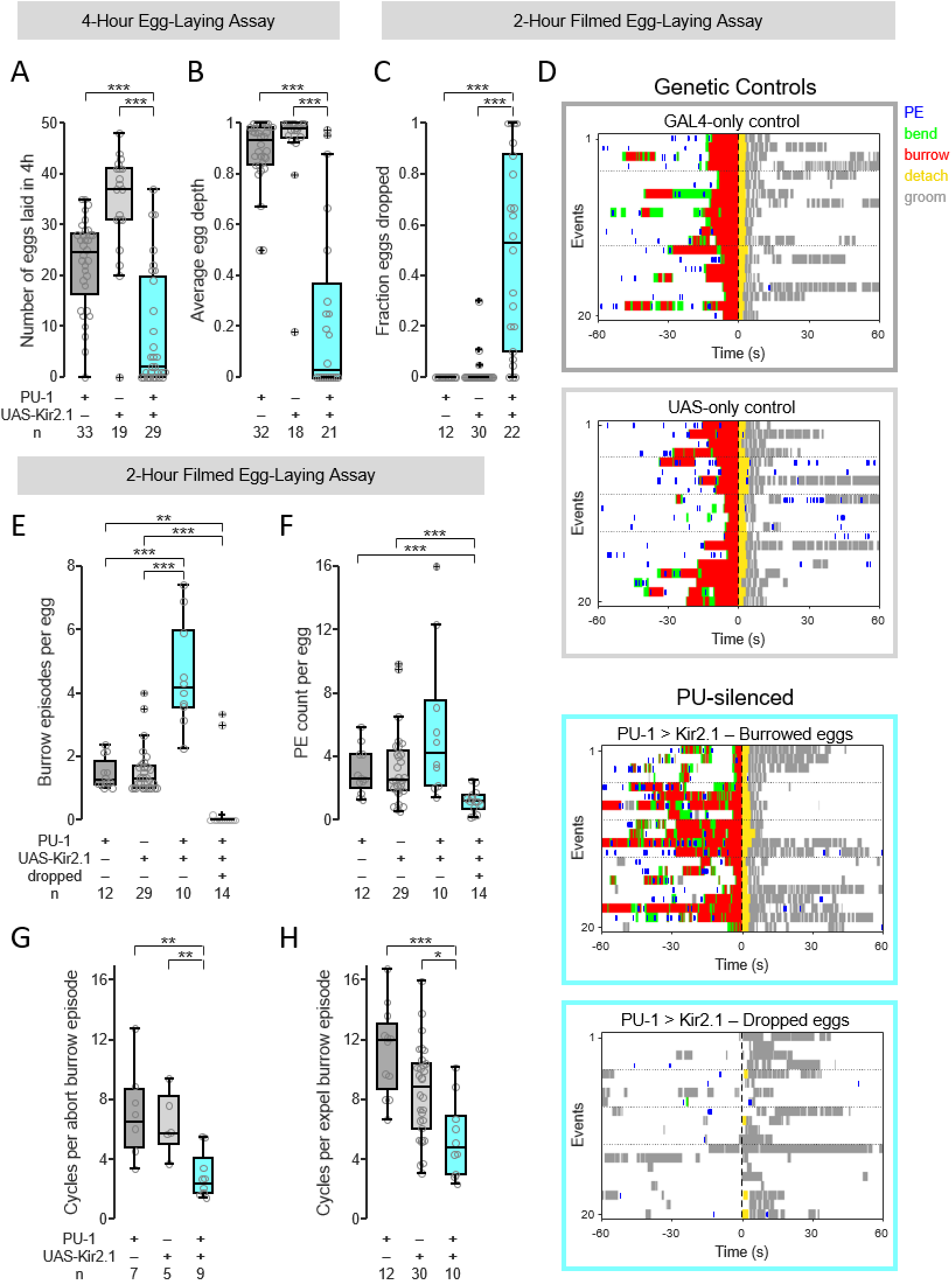
Silencing PU neurons reduces egg output and disrupts multiple components of the egg-laying sequence. (A) Number of eggs laid on 1 % agar in a 4-hour window. For all panels in this figure, GAL4-only control is PU-1 > myr-GFP and UAS-only control is empty-SplitGAL4 > Kir2.1. For all panels in this figure, *p<.05, **p<0.01, ***p<0.001, two-sided Wilcoxon rank sum test followed by Bonferroni correction. (B) Average depth of penetration of laid eggs (see Methods). (C) Percentage of eggs dropped without burrowing. Only flies that laid four or more eggs were considered for this analysis as well as in panels E-H. (D) Ethograms of egg-laying behavior for two genetic controls (top two panels), and for PU-silenced flies that burrowed eggs (third panel) or dropped eggs (bottom panel). Time = 0 marks the time of completed egg deposition (“egg out”). (E) Average number of burrowing episodes per egg-deposition event. Only flies that exhibited three or more burrowed eggs were considered for the first three groups, or three or more dropped eggs for the fourth group were considered here and in (F). (F) Average number of proboscis extension (“PE”) events preceding egg deposition. (G) Average number of cycles per aborted burrowing episode. Only flies that exhibited three or more aborted burrowing episodes were considered here. (H) Average number of cycles per egg-deposition burrowing episode. Only flies that exhibited three or more egg-deposition burrowing episodes were considered here.

We further examined the behavioral consequences of PU-silencing by filming flies in small egg-laying chambers. We observed that on average flies dropped 49% of the eggs without burrowing (Figures 4C and 4D, bottom panel), whereas the remaining 51% were deposited following a burrowing episode. PU-silenced flies that deposited eggs following burrowing exhibited a seven-fold increase in aborted burrowing episodes prior to egg deposition (PU-1>Kir2.1, 3.6 aborted episodes versus 0.5 for both genetic controls; Figures 4D, third panel, and 4E). As a consequence of the increase in aborted episodes, these flies exhibited an excess of exploration behavior (Figure 4F). Burrowing episodes were comprised of fewer cycles compared to episodes in control flies (Figures 4G and 4H). Moreover, burrowing episodes culminating in egg deposition resulted in the premature release of the egg on the substrate surface. These data suggest that PU senses the progression of the egg into the ovipositor during a burrowing episode, promoting persistent burrowing to achieve subterraneous egg deposition.

We observed a second distinct phenotype upon PU-silencing; silenced flies drop eggs without exhibiting burrowing or any of the other behavioral components that precede egg deposition. This phenotype was observed in 49% of egg-laying events and was exhibited by 19 out of 22 PU-silenced flies (Figure 4C). Dropped eggs were only rarely observed in control flies (0% in 12 GAL4-only control flies, and 2% in 3 of 30 UAS-only control flies; Figure 4C). Thus, the silencing of PU neurons resulted in deficits in exploration, burrowing, and possibly fertilization, suggesting that PU feedback is involved in the expression of multiple components of the behavioral sequence that surrounds egg deposition.

### PU activity determines timing and direction of transition from burrowing

We next explored the function of PU neurons by targeted expression of the red-light activated channelrhodopsin, CsChrimson ^44^. In initial experiments, we asked whether stimulation of PU neurons could elicit components of egg-laying behavior in flies lacking an egg in the uterus. We photo-stimulated either virgins or mated females with a genetic block to ovulation to assure that flies were unable to release eggs into the uterus ^37^(see Methods). Photo-stimulation did not elicit any of the components of egg-laying behavior in either group of flies.

We therefore devised a physiological paradigm in which we photo-stimulated PU neurons in the context of egg-laying. We have shown that PU neurons are activated upon passage of the egg from the uterus to the ovipositor and that their activity returns to baseline upon completed expulsion. We reasoned that prolonged PU neuron activation beyond completed egg deposition may mimic the continued presence of the egg within the ovipositor, and delay progression along the behavioral sequence. In control flies, burrowing ceases after egg deposition and the fly detaches from the egg, grooms its ovipositor, and transitions to the “reset” phase. We photo-stimulated PU neurons for various durations during burrowing, immediately prior to egg deposition (655nm at 8 μW/mm^2^, PU-1 > CsChrimson; Figure 5A). These flies deposit their eggs but prolonged PU neuron activation beyond deposition resulted in the aberrant persistence of the burrowing episode without transitioning to detachment and grooming (Figure 5B; compare with “no light” control in top plot). The duration of aberrant burrowing after egg deposition was correlated with the duration of photo-stimulation (r = 0.36, p<0.001). In 43% of the events, burrowing persisted throughout the duration of photo-stimulation and ceased upon light offset. In the remaining events burrowing persisted for variable durations but stopped prior to light offset (Figure 5B). These two behaviors may reflect the action of PU in the persistence and abortion observed during normal burrowing episodes.

**Figure 5.**
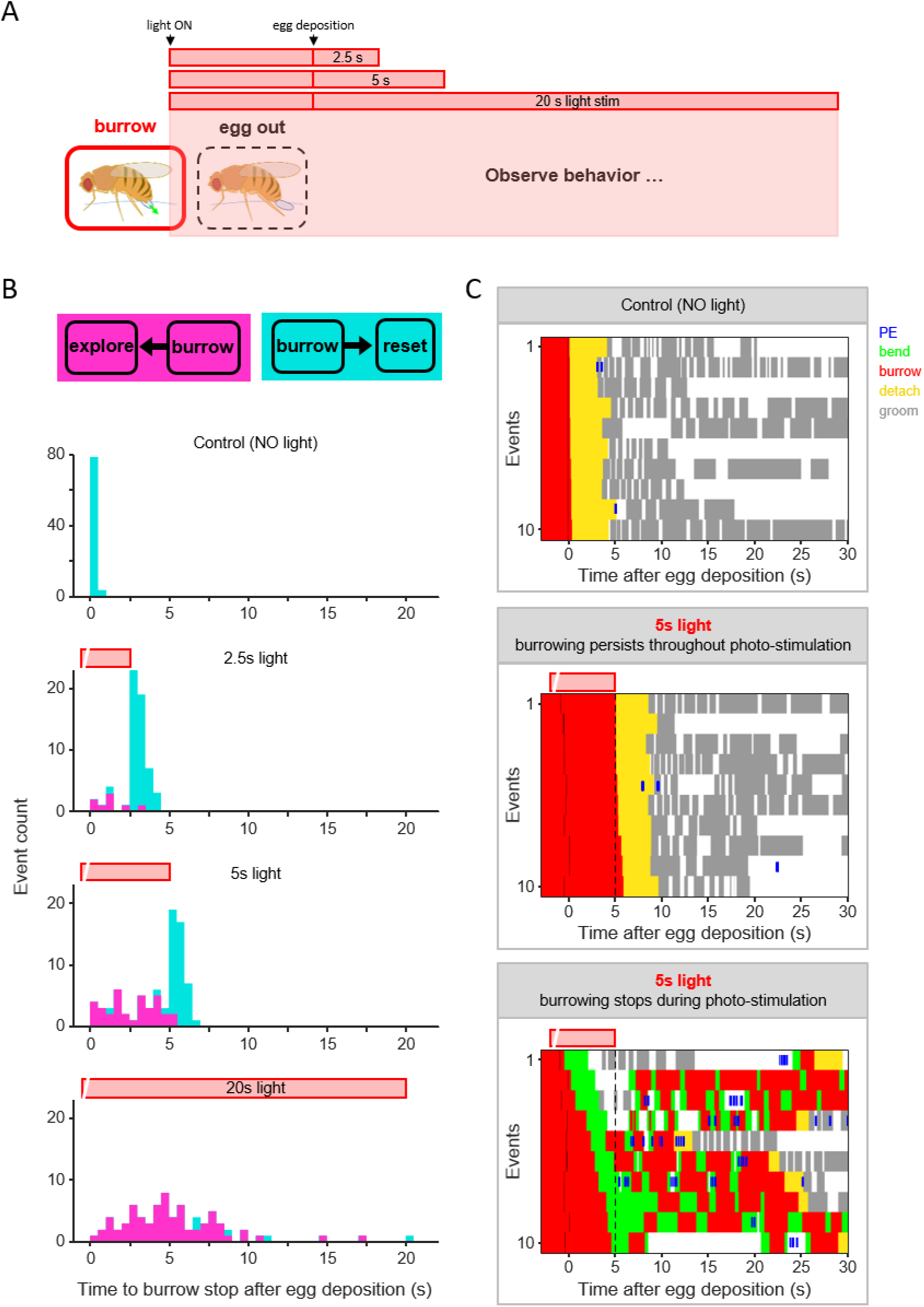
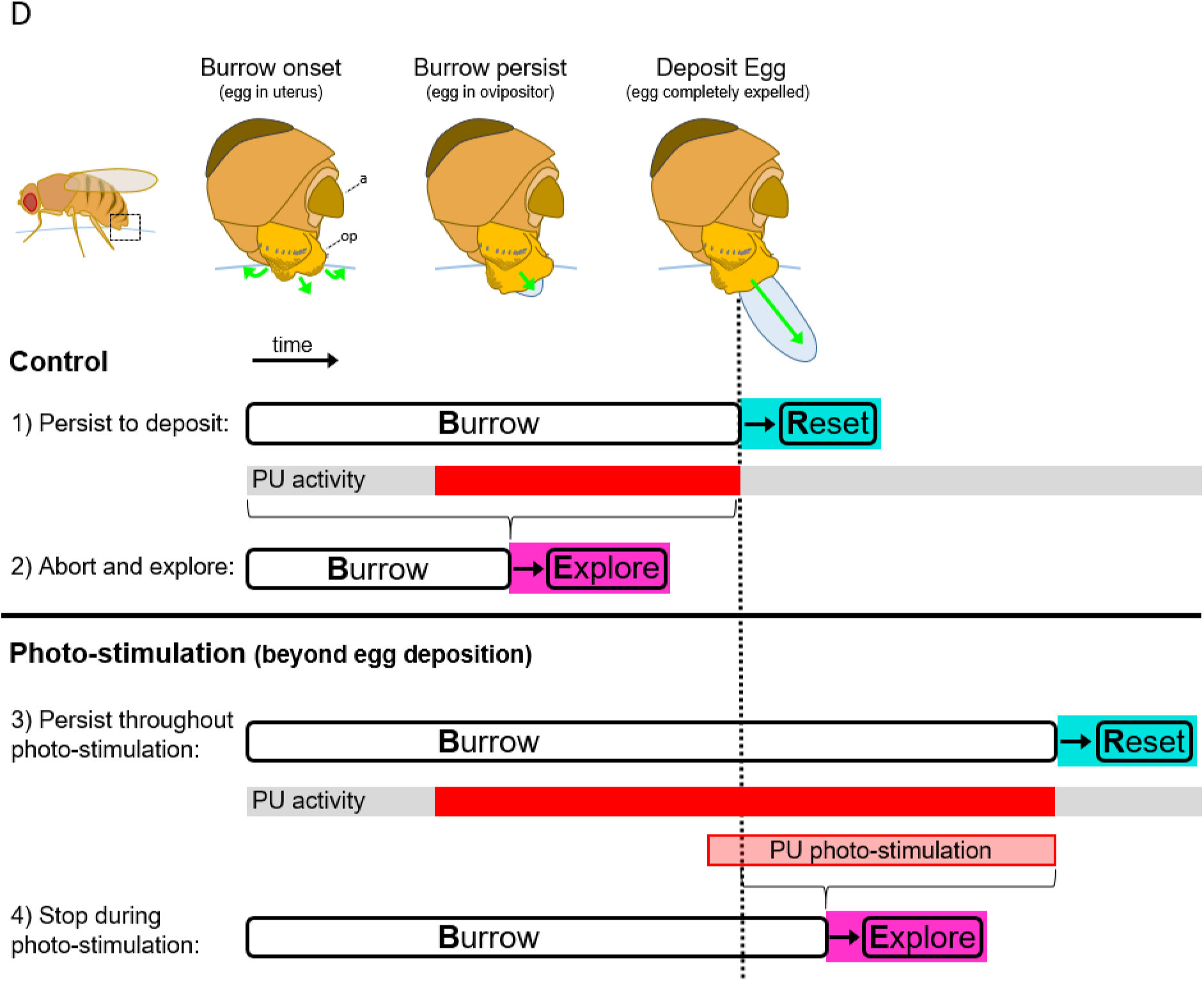
PU activity determines timing and direction of transition from burrowing. (A) Schematic of photo-stimulation paradigm. Photo-stimulation (655nm light at 8μW/mm^2^) was initiated during burrowing immediately prior to completed egg deposition and sustained for variable amounts of time after egg deposition (see Methods). (B) Stacked distributions of the timing that burrowing stopped after egg deposition for control (top) and stimulus conditions. Events are color coded according to which transition was made after burrowing stopped. Magenta, flies reverted to exploration; Cyan, flies progressed to the “reset” phase (see Methods and Figure S4C). The period of photo-stimulation is indicated by the red bar above each plot. Data presented represent a total of 298 events from 16 flies. (C) Behavioral annotations for no-light control events (top) and 5-second photo-stimulation events, separately depicting those where burrowing persisted until light off (middle) from those where burrowing was stopped prior to light off (bottom). The period of photo-stimulation is indicated by the red bar above each plot, and photo-stimulation offset by a vertical black dashed line. For each stimulation event, the black tick mark near time = 0s indicates the timing of completed egg deposition. (D) Proposed model for how PU activity determines the timing and direction of transitions made from burrowing. Top, control behavior; bottom, behavior during photo-stimulation. The box shown on the fly (top left) indicates the region depicted in higher detail to the right (‘a’, anal plates; ‘op’, ovipositor). The vertical black dashed line indicates the timing of completed egg deposition (“egg out”). PU neurons are at baseline (gray) at the onset of burrowing, become activated (red) upon progression of the egg into the ovipositor during burrowing, and return to baseline upon completed egg deposition. For both condition 2 (control) and condition 4 (photo-stimulation), termination of burrowing at any time within the bracketed region results in the reversion to exploration.

Flies that continue to burrow throughout photo-stimulation abruptly stopped burrowing upon light offset and transitioned to the behavioral sequence normally triggered by egg deposition (Figure 5C, middle panel; compare with “no light” control in top panel; Figure S4A). This phenotype was observed for all three stimulus durations (51/51 for 2.5s stim, 41/41 for 5s stim, and 1/1 for 20s stim; Figure 5B; cyan bars indicate events that transition to “reset”; Figure S4B). These data support the argument that PU neurons are activated by the progression of the egg into the ovipositor and drive the persistence of burrowing. In addition, these observations suggest that a decrease in PU activity signals the completion of egg deposition, resulting in the transition to detachment, grooming, and the “reset” phase (Figure 5D).

In 57% of the photo-stimulation events burrowing persisted for variable durations but stopped prior to light offset. In these events, flies reverted to exploration after burrowing stopped rather than transitioning to the post-deposition sequence (8/9 for 2.5s stim, 35/39 for 5s stim, and 70/74 for 20s stim; Figure 5B; magenta bars indicate events that revert to exploration; Figures 5C, bottom plot, S4B and S4C). Although these flies have deposited an egg they explore and then burrow in new locations. This phenotype appears to recapitulate the behavior observed after aborting a burrowing episode in normal egg-laying behavior. During a burrowing episode in control flies, prolonged PU activation without deposition may signal the inability to deposit an egg and result in abortion of the episode (Figure 5D). In the photo-stimulation experiment, the flies may be unaware of having laid an egg, and the prolonged activation of PU neurons may signal that egg deposition is unlikely, and the fly aborts the episode.

These flies persisted in exploration and exhibited numerous burrowing episodes for up to several minutes beyond light offset despite the absence of an egg in the uterus (6.4 ± 6.1 episodes, last episode initiated 24s ± 38s after light offset; mean ± SD; Figures S4D and S4E). After the decision to abort and explore, a decrease in PU activity (photo-stimulation offset) no longer triggers the transition to “reset.” Thus, a decrease in PU neuron activity only signals the completion of egg deposition and the transition to post-deposition behaviors in the context of an ongoing burrowing episode. Sensory information obtained during a component action therefore governs the progression along the behavioral sequence.

## DISCUSSION

We have characterized the structure of egg-laying behavior in the fly and demonstrate that it consists of a sequence of component actions analogous to Nikolaas Tinbergen’s “reaction chain”^1^. Tinbergen portrayed innate behaviors as a “reaction chain” in which each component action of the sequence enhances the probability of encountering “sign stimuli” that promote progression to a subsequent component. One classic example provided by Tinbergen is the mating ritual of stickleback fish. During the mating season a gravid female approaches a male that displays a red underbelly, prompting the male to perform a zig-zag dance. The dance entices the female to approach closer and the male then leads the female to the nest. The male grazes her, promoting spawning, and then releases sperm upon the clutch. The organization of component actions into a “reaction chain” provides decision points at the junctures of component behaviors that ensure the successful progression towards the consummate act that satisfies the drive.

Egg-laying behavior in a fly is a repeating cycle of component behaviors that transitions between exploration, deposition, and a “reset” phase that includes ovulation ^14,18^. Our data suggests that the individual components of egg-laying behavior are not simply motor acts but also acts of sensory evaluation that shape the motor program. During early exploration, substrate cues are encountered upon walking and proboscis sampling that may identify a suitable location for egg deposition ^10,13^. The flies then initiate more refined, local exploration involving abdominal bending to permit ovipositor sampling ^45^. This behavior may provide sensory cues that promote the transition to burrowing. During burrowing the ovipositor is used to score the surface, extend into the substrate and deposit the egg subterraneously. The deposition of the egg triggers egg detachment and the transition to the final behavioral phase, grooming and ovulation, facilitating the re-initiation of the sequence. We have identified a cluster of sensory neurons (PU neurons) that signal the progression of the egg through the ovipositor during burrowing and may facilitate the transition to the final “reset” phase. Behavioral analysis along with genetic manipulation suggests that information from the PU neurons can either drive the persistence of burrowing resulting in egg deposition, or prompt the cessation of burrowing if the egg cannot be expelled. Finally, diminished activity in these neurons during burrowing signals the completion of egg expulsion and may initiate the transition to the “reset” phase. These results provide a logic for a “reaction chain” in which sensory information at critical junctures allows flexible adjustments in component behaviors to achieve subterraneous egg deposition.

Structuring an innate behavior in this manner provides decision points at the junctures of the individual component actions. The presence of such decision points affords the organism with flexibility that permits adjustment to accommodate variability in the internal or external environment. The organization of innate behaviors into a sequence of component actions may confer additional adaptive advantages. Individual components in an innate behavioral sequence may be differentially responsive to the distinct sensory stimuli that promote transitions along the sequence. A given stimulus may behaviorally impact only one of the components in the sequence. Each component therefore establishes a context that filters relevant sensory input. For example, during the hunting of bees, digger wasps first visually identify a target bee. The odor of bees has no impact during this visual search but once a bee has been spotted, the bee odor triggers an acute strike ^1,46^. Similarly, we observe context dependent behavioral responses to the activation of PU neurons. Photo-stimulation of PU during a burrowing episode results in persistent burrowing, whereas activation at other times in the sequence does not elicit a behavioral response. Moreover, only during burrowing does a decline in PU activity result in the transition to the “reset” phase. Thus, the behavioral impact of sensory stimuli differs for each of the components in a sequence. Each component therefore displays selective attention to distinct stimuli that structures the transition between behaviors to accommodate a complex and variable sensory environment.

In addition, each component in the sequence affords an entry point for adaptive evolutionary change. Changes in specific components of egg-laying behavior that accommodate a new ecological niche can occur without perturbing the overall sequence. For example, changes in substrate preferences during early exploration may occur as an evolutionary adaptation to a changing environment ^9,11,47^. Alterations in subsequent components such as burrowing may then be necessary to accommodate changes in the properties of the novel substrate. For example, *D. melanogaster* and *D. suzukii* have different preferences for the site of egg-laying ^9^. *Suzukii* females prefer to deposit eggs within firmer, ripe fruit, whereas *melanogaster* favors softer, rotting fruit. Egg-laying behavior is comprised of the same sequence of component actions in the two species, but *suzukii* females exhibit dramatically prolonged burrowing episodes ^17^. Episodes in *suzukii* can persist for over 100 seconds, whereas *melanogaster* burrowing on the same substrate does not extend for more than 9 seconds. Interestingly, *suzukii* have also evolved an enlarged and serrated ovipositor ^48^. These changes do not alter the sequence of behaviors, but illustrate evolutionary adaptations that allow *suzukii* to deposit eggs within firmer fruits. A similar logic holds for male *Drosophila* courtship behavior, where adjustments in different steps in a conserved sequence (e.g. foreleg pheromone sampling, singing) can be independently altered and these modifications play critical roles in sexual isolation between related species ^49–51^.

An innate behavioral repertoire is thought to be initiated by higher order brain centers that represent a specific motivational state, or drive ^1,19–22,52,53^. These centers are activated by stimuli relevant to the drive and then select an appropriate motor program for action ^16,23,24,54^. A signal is then transmitted to pre-configured circuits in the ventral nerve cord or spinal cord that are capable of producing a coordinated sequence of motor actions ^25–28^. Pivotal intermediaries in this pathway are the descending interneurons that link the output of higher brain centers with the appropriate local circuits in the ventral nerve cord ^16,23,54–60^. One intermediary eliciting components of egg-laying in *D. melanogaster* has been recently identified, the descending oviDN cluster of neurons ^16^. OviDNs are necessary and causal for the expression of multiple components of the egg-laying sequence: abdominal bending, burrowing, and egg expulsion. Higher order brain centers disinhibit the oviDN cluster following mating and modulate oviDN activity in response to mechanical and gustatory stimuli presented to the legs. Thus, oviDNs are poised to induce the transition from early exploration (walking and proboscis sampling) to later steps in the sequence resulting in egg deposition. Although activation of oviDN is capable of eliciting the sequence from abdominal bending to egg expulsion, our observations suggest transitions along this late sequence are exquisitely sensitive to ongoing sensory feedback. We observe that bending and ovipositor sampling do not always lead to burrowing, suggesting that this transition is regulated by feedback from ovipositor sensory neurons. Furthermore, the duration of burrowing and the transition to the “reset” phase are informed by feedback from PU neurons that sense the progression of the egg in the ovipositor. Thus, once oviDN is selected for action by higher brain centers, sensory information acquired during the behavioral sequence governs the progression to satisfy the drive.

## ACKNOWLEDGEMENTS

We thank David Anderson, Cory Root, and Larry Abbot for comments on the manuscript; Daisuke Hattori, Barbara Noro, Jenna Browning-Kamins, Juliana Remark, Ariel Pourmorady, Rares Mosneanu, and Caroline Haoud for technical assistance; Nicolas Gompel, Ellen Lumpkin, Angelica Vina-Albarracin, Kenny Kay, Lucas Tian, and members of the Axel and Dickson laboratories for discussions; Gerry Rubin, Kristin Scott, Daisuke Hattori, Barry Dickson, and David Anderson for fly stocks; and Clay Eccard for assistance in preparation of the manuscript. This work was supported by the Simons Foundation (R.A.) and the Howard Hughes Medical Institute (R.A.). R.A. is an HHMI investigator.

## AUTHOR CONTRIBUTIONS

K.M.C. designed the study, built all experimental devices, and performed all experiments and analyses. K.M.C. and R.A. wrote the manuscript.

## METHODS

### EXPERIMENTAL MODEL AND SUBJECT DETAILS

#### Flies

The G4DBD split components of PU-1 and PU-2 in *attp2* were generated as previously described 3^9,61^. The p65AD split component of PU-1 and PU-2 (in *attp40*), the PU parent LexA lines (all in *attp40*), *GMR23C03-LexA, empty-SplitGAL4* (*pBPp65ADZp* in *attP40*; *pBPZpGDBD* in *attP2*), *LexAop-mCD8-GFP* in *attp2*, and *UAS-GCamp6f* in *attp40* were obtained from Bloomington Drosophila Stock Center. *UAS-mCD8-GFP* in *VK00005, UAS-tdTomato* in *VK00005*, and *UAS-myr-GFP* in *VK00005* were gifts from Gerry Rubin. *hs-FLP* was a gift from Kristin Scott ^62^. *UAS-KIR-tdTomato* in *VK00005* was a gift from Daisuke Hattori. *UAS*-(*FRT.stop*)*CsChrimson* in *attp18, LexAop-Kir2.1*^63^ and Canton-S were gifts from Barry Dickson. *UAS-CsChrimson-tdTomato* in *VK00005*^44,64^ was a gift from David Anderson.

### METHODS DETAILS

#### Wild-type and Loss-of-Function Behavior

Flies were reared at 25°C and 55% relative humidity on a 12-hour light: 12-hour dark photoperiod in vials containing standard cornmeal-agar food. With the exception of experiments using virgins, females used in egg-laying experiments were genotyped under CO_2_ anesthesia within one day of eclosion and transferred to a vial containing an enriched medium (Nutri-Fly GF, Genesci Scientific) in a ratio of four females to five males, with a minimum of 12 and a maximum of 20 females per vial ^37^. Egg laying experiments were performed five to seven days-post-eclosion. All experiments were initiated ± 2 hours of lights-off and performed within a temperature and humidity controlled room.

For the characterization of the time interval between eggs and speed surrounding egg deposition (Figures S1A and S1B), single females were filmed in parallel within a laser-cut acrylic assembly containing four chambers (10 mm x 18 mm x 16 mm). Substrate comprised of 1% agarose (Affymetrix Agarose-LE, 32802) and 5% acetic acid (volume:volume; Sigma-Aldrich 338826) was poured into each chamber and allowed to set for 30 minutes before females were introduced by gentle aspiration ^13,37^. The assembly was placed atop a red led panel light (Advance Illumination) for illumination, and adjacent to a mirror positioned at a 45° angle, allowing for the flies to be simultaneously filmed from the top and side perspective. Video recording was performed using a USB3 camera (FL3-U3-13Y3M-C, Point Grey) attached to a 6x macro zoom lens (Edmund Optics #68-667) at 2 Hz (262 x 445 x 390 pixels per chamber). Experiments lasted three hours.

For quantifying behavior in high resolution (Figures 1, 2A-C, 2F, 2G, and 4C-H; “2-Hour Filmed Egg-Laying Assay”), single females were filmed in parallel within a custom-designed 3D-printed assembly containing six chambers (Shapeways). Individual chambers were 4.1 mm deep that tapered from top to bottom (from 7.3 mm × 5.8 mm to 6.7 mm × 4.3 mm). The bottom face of the chamber was painted black to facilitate visualization of the fly and eggs. One side of the chamber was open to a reservoir within which the agar-based substrate plus 3% acetic acid was poured and allowed to set for 30 minutes. Flies were introduced by gentle aspiration one at a time through a loading hole in an acrylic window that was slid into place along grooves at the top of the assembly, enclosing the fly within the chamber and allowing for viewing. The assembly was placed at the center of a 2-inch off-axis ring light for illumination (530nm, Metaphase Technologies) and video recording was performed from the top down using a GigE camera (Basler Ace acA2000-50gm-NIR) attached to a 0.5X telecentric lens (Edmund Optics #54-798) at 20 Hz (682 x 540 pixels per chamber). Flies from experimental and genetic control groups were run simultaneously in neighboring chambers. Experiments lasted two hours.

For the study of egg depth of penetration across substrates of varied hardness (Figures 2D, 2E, 4A, 4B, S3A, and S3B; “4-Hour Egg-Laying Assay”), we developed a laser-cut acrylic assembly with 16 individual chambers (18.5 mm x 18.5 mm x 6 mm). The assembly was covered on the top and open on the bottom, and had a narrow slot in the middle to allow a thin plastic barrier to be slid into place. Flies were introduced by gentle aspiration through loading holes on the top which were then sealed with tape to enclose the chamber. The flies were allowed to habituate to the chamber while the substrate was prepared. 40 ml of substrates containing agarose and 3% acetic acid was poured into the lid or base of a 120 mm square petri dish (Greiner) and allowed to set for 30 minutes. The assembly was then placed atop the substrate. The experiment was initiated by removal of the thin plastic barrier separating the flies from the substrate, and the whole assembly was then placed in a dark enclosure for four hours. Experiments for all agarose concentrations were run simultaneously. At the end of the experiment, the flies were removed and the eggs per chamber were counted and scored for depth of penetration. An egg was given a score of 1 if it was deposited entirely beneath the substrate surface, with only the egg’s spiracles (breathing tubes) exposed. If only part of the egg was beneath the surface, it was given a score of 0.5, whereas if it was entirely resting on the surface it was scored as 0.

For the assessment of virgin receptivity and egg-laying after a single mating (Figures S3C-E; “Single-mating, 48-hour egg totals”), experiments were performed similarly as described before ^63^. Receptivity was scored over a one-hour period. For egg-laying, after mating, females were transferred to small chambers (9.5 mm x 8 mm x 10 mm) containing cornmeal-agar-molasses food and again 24 hours later. Eggs were counted at the 24- and 48-hour mark, and combined. The number of unhatched eggs was counted 48 hours after flies were removed.

#### Immunostaining and Confocal Microscopy

Flies were anesthetized with CO_2_ and minimally dissected (decapitated, and with small incisions made in the dorsal thorax and anterior abdomen) before being fixed with 2% PFA/PBL (2% paraformaldehyde in 75mM lysine, 37mM sodium phosphate buffer, pH7.4) for 2 hours at room temperature (RT). The flies were then removed, and the brains, ventral neve cord, and lower reproductive tract were dissected in 1xPBS, blocked with 10% normal goat serum diluted in PBS containing 0.3% Triton X-100 (PBST) for 30 minutes at RT, and then incubated in a primary antibody mix overnight at 4°C. Subsequently, the tissue was washed for multiple rounds with PBST before being incubated with a secondary antibody mix overnight at 4°C. A final round of PBST washing occurred before the tissue was mounted using VectaShield (Vector Laboratories) and imaged using a LSM 710 laser scanning confocal microscope using a 25x/ 0,8 DIC or 40x/ 1,2 W objective (Zeiss). Primary antibodies used were mouse anti-bruchpilot (nc82, 1:10, Developmental Studies Hybridoma Bank), chicken anti-GFP (1:1000, Aves Labs), and rabbit anti-DsRed (1:500, Clontech). Secondary antibodies used were Alexa Fluor 633 goat anti-mouse, Alexa Fluor 488 goat anti-chicken, and Alexa Fluor 568 goat anti-rabbit (all at 1:200, Life Technologies). To visualize muscle fibers in lower reproductive tracts, Alexa Fluor 633 phalloidin was included in the secondary antibody mix (1:200, Life Technologies). Acquired images were processed using the Fiji distribution of ImageJ (NIH).

#### Thoracic Dissection for Calcium Imaging

Females were genotyped under CO_2_ anesthesia within one day of eclosion and housed in a vial containing standard cornmeal-agar food in a 1:1 ratio with males, and a maximum of 10 females per vial. Females were selected for experimentation between 3 and 20 days-post-eclosion. For an experiment, a female was anesthetized on ice and its wings were removed with fine Vannas scissors before being mounted, ventral-side up, on a square acrylic platform using a small drop of UV-curable glue (AA 3104, Loctite) and UV illumination (LED-200, Electro-Lite Co. Bethel, CT USA). During curing, the head and abdomen were lightly pressed down to ensure complete mounting from the head to just anterior of the anal plates. All legs were cut short at the trochanter before, using a 30-gauge needle, a thin well was created with petroleum jelly that encompassed the remaining leg coxa and ranged from the neck connective to the anterior abdomen. A custom-designed imaging platform, with a hole (1 mm x 750 μm) at the bottom of a pyramidal basin was positioned using putty such that the hole was centered on the hind leg coxa. The basin was filled with external saline (108 mM NaCl, 5 mM KCl, 2 mM CaCl_2_, 8.2 mM MgCl_2_, 4 mM NaHCO_3_, 1 mM NaH_2_PO_4_, 5 mM trehalose, 10 mM sucrose, 5 mM HEPES pH 7.5, osmolarity adjusted to 275 mOsm), before fine forceps were used to remove the remaining coxa of the middle and rear legs, along with the surrounding preepisternum, the internal sternal apophysis, and any visible trachea, revealing an rectangular window above the abdominal ganglion and proximal extent of the main abdominal nerve. Finally, to clear off debris, the basin was drained with a Kimwipe, and fresh saline was gently flushed over the window.

#### Calcium Imaging During Egg Expulsion

Experiments were initiated immediately upon completion of the dissection, and flies were typically imaged for 30-60 min. The acrylic platform was secured with a filter holder (Thorlabs DH1) positioned adjacent to a camera and high magnification lens setup (Point Grey USB3 camera, CM3-U3-13S2M-CS; InfiniProbe S-80 Right Angle Video Microscope lens) and band-pass filter (Thorlabs FGB25S) that, when illuminated by a nearby infrared (850nm) LED lamp, allowed for high resolution video recordings of the posterior abdomen concurrent with 2-photon imaging. Only flies that harbored an egg in the uterus were used for experiments.

2-photon experiments were performed using an Ultra microscope (Bruker) coupled to a Ti:Sapphire laser (Chameleon Vision, Coherent), with a GaAsp detector (Hamamatsu Photonics) for Gcamp6 and a photomultipier tube for tdTomato imaging. We used a 40x/0.80 NA water-immersion objective (Nikon), and the laser was tuned to 925 nm; the power measured after the objective ranged from 5-7 mW. The abdominal ganglion and main abdominal nerve were located using the microscope oculars and positioned near the center of the field-of-view by 2-photon imaging. Using the tdTomato anatomical marker, a stretch of the main abdominal nerve where the axons were separated and running in parallel was selected for the coronal section ^42^. Coronal section imaging was performed at 10 Hz, covering 42.4 μm in X and 60 μm in Z (512 by 85 pixels per image; 1.2 μs pixel dwell time). Small adjustments in the X and Z dimensions were made as needed throughout the experiment to compensate for drift.

#### Optogenetic Stimulation During Behavior

Flies were reared and collected as described above with two exceptions: 1) flies were reared and maintained in the dark, and 2) all trans-Retinal (0.4mM, Santa Cruz Biotechnology) was included in the enriched medium. The high-resolution filming apparatus described above was slightly modified, as the flies were instead illuminated with an infrared 2-inch off-axis ring light (880nm, Metaphase Technologies), a single 655nm high power LED (Luxeon Star) was installed adjacent to the video lens to deliver red light stimulus, and a band-pass filter was mounted in front of the lens. A custom MATLAB graphical user interface (GUI) was used to select the stimulus condition and control the timing and intensity of the light stimulus via an Arduino UNO (Arduino) and LED controller (BuckPuck 700mA, Luxeon Star). As the egg neared completed deposition, a trigger was pressed that turned the light stimulus on. The instant the egg was fully deposited, a second trigger was pressed, initiating a countdown timer whose duration was determined by the selected stimulus condition, which ultimately turned the light off upon reaching t = 0. Only one fly was experimented with at a time, and the precise timing of the light stimulus was extracted from a narrow portion of an adjacent chamber that was not blocked by the long-pass filter. Individual flies contributed a minimum of 15 events (5 events each for control and two experimental conditions) and a maximum of 20 events (five events each for control and all three experimental conditions) to the final data set.

The photo-stimulation intensity used in final experiments (8 μW/mm^2^) was determined as the minimum intensity that reliably produced either phenotype in the 5s stimulation condition (35/73 at 2μW/mm^2^, n = 15 flies, 61/65 at 4μW/mm^2^, n = 14, and 76/80 at 8μW/mm^2^, n = 16). Phenotypes were reliably observed using the second PU split-Gal4 line in response to 5s stimulation at 8 μW/mm^2^ (PU-2>CsChrimson, 60/60, n = 12). Genetic controls did not exhibit a phenotype in response to 5s stimulation at 8 μW/mm^2^ (PU-1 > myr-GFP, 0/50, n = 10 flies; empty-SplitGAL4 > CsChrimson, 0/50, n = 10).

For the experiments performed in virgins, PU-1 X UAS-CsChrimson crosses were set up on all trans-Retinal, virgins were isolated from males upon eclosion, and experiments were performed at 1 day-post-eclosion. To explore the consequences of photo-stimulation in mated females with a genetic block to ovulation, experimental flies included *GMR23C03-LexA* and LexAop-Kir2.1. GMR23C03-LexA drives expression within previously identified oviduct motor neurons ^37^ that, when silenced, prevent eggs from passing through the oviduct (K.M.C., unpublished data).

### QUANTIFICATION AND STATISTICAL ANALYSIS

#### Wild-type and Loss-of-Function Behavior Data Analysis

The first step in all video behavioral analysis was to record the timing of deposition for all eggs, which served as a landmark for further analysis. For the high-resolution assay, the video was segmented in one of two ways. Most data were analyzed over ± 60 seconds of egg deposition, with the exception of the analysis of burrowing behavior across varied substrate hardness (Figures 2F and 2G). Here, behavior was analyzed over a contiguous video segment spanning three to eight egg-laying events, beginning from just after the first egg was deposited.

Segmented videos were then manually annotated frame-by-frame in a custom MATLAB GUI. The start and end frame of each behavioral event was recorded, with the exception of “PE,” where only a single frame was noted at the onset of each proboscis contact. “Egg out” was determined as a single frame at the completion of egg deposition. All varieties of grooming behavior were scored as “groom.” As abdominal bending is obligate for the induction of burrowing, all burrowing events were preceded by “bend,” and also followed by “bend” if not resulting in egg deposition. The timing and count of burrow cycles were determined by observing individual episodes in real-time. All burrowing episodes were analyzed, including those occurring on the rigid chamber walls, except for the analyses performed across varied substrate hardness (Figures 2F and 2G) and PU-silenced behavior (Figures 4E-H) which were limited to burrowing episodes occurring on the substrate.

In the lower resolution assay involving simultaneous top- and side-views (Figures S1A and S1B), the speed surrounding egg deposition was determined by comparing the distance between the fly’s center-of-mass in all three dimensions across successive frames (500 ms). For the high resolution assay, DeepLabCut ^65^(DLC, a feature detection algorithm) was used to track the X-Y position of the posterior tip of the scutellum on the thorax, and speed was estimated by comparing the distance between this position across 10 frames (500 ms). In rare cases where the scutellum was not in view, the position of the proboscis was used instead. In a given frame, a fly was considered to be moving if its speed, smoothed by a 1-second moving average, was greater than a threshold of 0.29 mm/s. This threshold was determined as the midline boundary between distributions of a two component Gaussian mixture model of all smoothed speed measures.

The Euclidean distance was calculated as the instantaneous distance between 5-second smoothed behavioral trajectories. Before smoothing, every instant for a given event was characterized by a 6-dimensional binary vector comprised of the five annotated behaviors and movement.

For the construction of the transition diagram (Figure 1F), a transition was defined as the subsequent component behavior to occur after the conclusion of the first component behavior. “PE” was only considered when expressed at distinct locations, spaced greater than 500 μm apart. With the exception of “PE,” behaviors were mutually exclusive. “PE” events that occurred during other behaviors were omitted from this analysis. Transitions were determined separately for behaviors happening before and after egg deposition. “PE” and “groom” occur both before and after egg deposition, though are only depicted once, according to where behavioral expression was highest. Transitions not shown correspond to these omitted behaviors and were of low probability (<0.07).

#### Calcium Imaging Data Analysis

To determine GCamp fluorescence levels in PU axons, the imaging data was first segmented into cell-specific Regions of Interest (ROIs). The location and shape of ROIs corresponding to all labelled axons across all frames was determined from the tdTomato channel using a semi-automated pipeline. The coronal sections of the MAN exhibited significant motion artifacts owing in part to motility of the immediately adjacent gut, as well as in response to abdominal muscle activity. This motion produced changes in the global location of the nerve within the field of view as well as local changes in the shape and relative location of individual axons.

We employed a ROI segmentation pipeline that relied on DLC predictions combined with custom MATLAB code. The number and relative location of individual axons was noted at the time of the experiment. We used DLC to track the center position of all identified axons, appearing as ellipsoids, in the tdTomato image stack. The DLC network was trained with a manually labelled set of diverse frames spanning the entire experiment. Additional frames were labelled as needed to produce accurate predictions (40-110 frames per experiment were labelled in total). DLC predictions were then used to select foreground ROIs from a binary thresholded image stack. If a given ROI included multiple predictions, such as when two axons were in close proximity, the ROI was divided according to distance from the DLC centers. Small ROIs were dilated to a minimum size of 30 pixels, and in frames where no ROI was found because of a dim tdTomato signal, a ROI representative of the average shape, centered on the DLC prediction, was selected. The raw fluorescence, F, was then calculated as the mean pixel value within the ROI bounds for each frame. ROIs corresponding to PU axons were determined by 2-photon imaging at the conclusion of the experiment. Axon projections were traced anteriorly, identifying PU axons as those that terminate in the ventral aspect of the abdominal ganglion. The raw fluorescence was converted to ΔF/F_0_ using a baseline determined as the median fluorescence value from recording onset up until 20 seconds prior to egg deposition, excluding ± 20 seconds surrounding ovipositor extrusion events. Example ΔF/F_0_ traces shown in figures and videos were smoothed by a 3-point moving average.

The timing of egg expulsion events and ovipositor extrusion events was manually annotated from the behavioral videos. The onset timing of incomplete egg expulsion or ovipositor extrusion in flies lacking an egg was defined as the first frame of maximal ovipositor extrusion; additional events were not scored until the ovipositor was fully retracted. The onset timing of complete egg expulsion was defined as the frame in which the posterior aspect of the egg first emerged from the ovipositor. Behavioral image acquisition was synchronized to 2-photon frame acquisition via a TTL-pulse coinciding with frame-offset.

To compare response magnitude across events and flies, the fluorescence data was integrated and normalized as follows. The integration window for both egg expulsion and ovipositor extrusion events was defined as t = 0 to t = 3 seconds after event onset. For comparing incomplete with complete egg expulsion (Figure 3F), the baseline, “0” value was determined as the median 3-second integral over the first contiguous stretch of 60 seconds leading up complete egg expulsion, excluding ± 20 seconds surrounding any egg expulsion event. The post-expulsion value was determined as the median 3-second integral from t = 10 to t = 20 seconds after complete egg expulsion. The maximum, “1” value for normalization was the maximum 3-second integral observed across all windows (baseline, incomplete expulsion, complete expulsion, and post-expulsion). For all neurons, this maximum was observed either in the incomplete or complete egg expulsion windows. For comparing incomplete egg expulsion with post-expulsion ovipositor extrusion events (Figure S2H), the minimum, “0” value was determined as the median of the first 60 (non-contiguous) seconds starting 10 seconds post egg-expulsion and excluding ± 20 seconds surrounding ovipositor extrusion events. The maximum value was determined as described above. For flies that expressed multiple events, the mean was used in plots and all analyses. Calcium imaging was performed using both PU-1 and PU-2 split-GAL4 lines, and the data was combined.

#### Optogenetic Behavior Data Analysis

For every egg-laying event, the timing of completed egg deposition and burrow termination was manually annotated frame-by-frame in a custom MATLAB GUI. The time of egg deposition was defined as the first frame where the egg reached maximum depth within the substrate. Burrow termination was defined as the frame associated with the onset of ovipositor detachment or lifting. The transition to the “reset” phase was determined if no additional burrowing episode occurred within 65 seconds of egg deposition. If additional burrowing episodes did occur, the onset timing of the last burrowing episode before “reset” was similarly determined as the last burrowing episode to precede a 65 second window free of burrowing. For the examples shown in Figure 5C, behaviors were manually annotated frame-by-frame as described earlier.

### SUPPLEMENTAL INFORMATION

**Figure S1.**
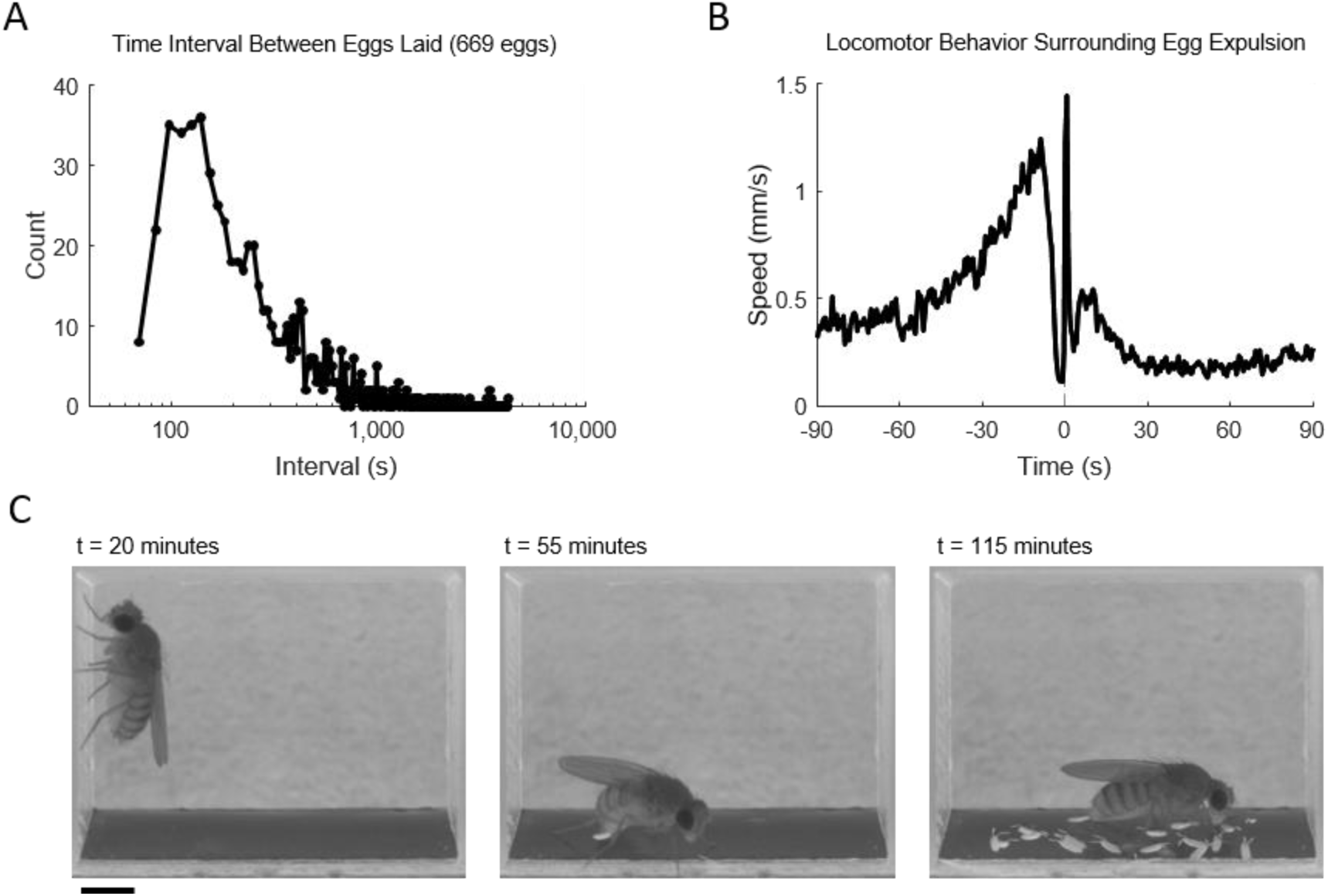
(A) The interval between successive egg-laying events observed over a 3-hour window. n = 669 eggs from 32 flies. (B) The average speed of flies over a 180-second window surrounding egg deposition. Time = 0 marks the time of completed egg deposition (“egg out”). (C) Example video snapshots of a single chamber from the high-resolution filming assay at three time points. Scale bar is 1 mm.

**Figure S2.**
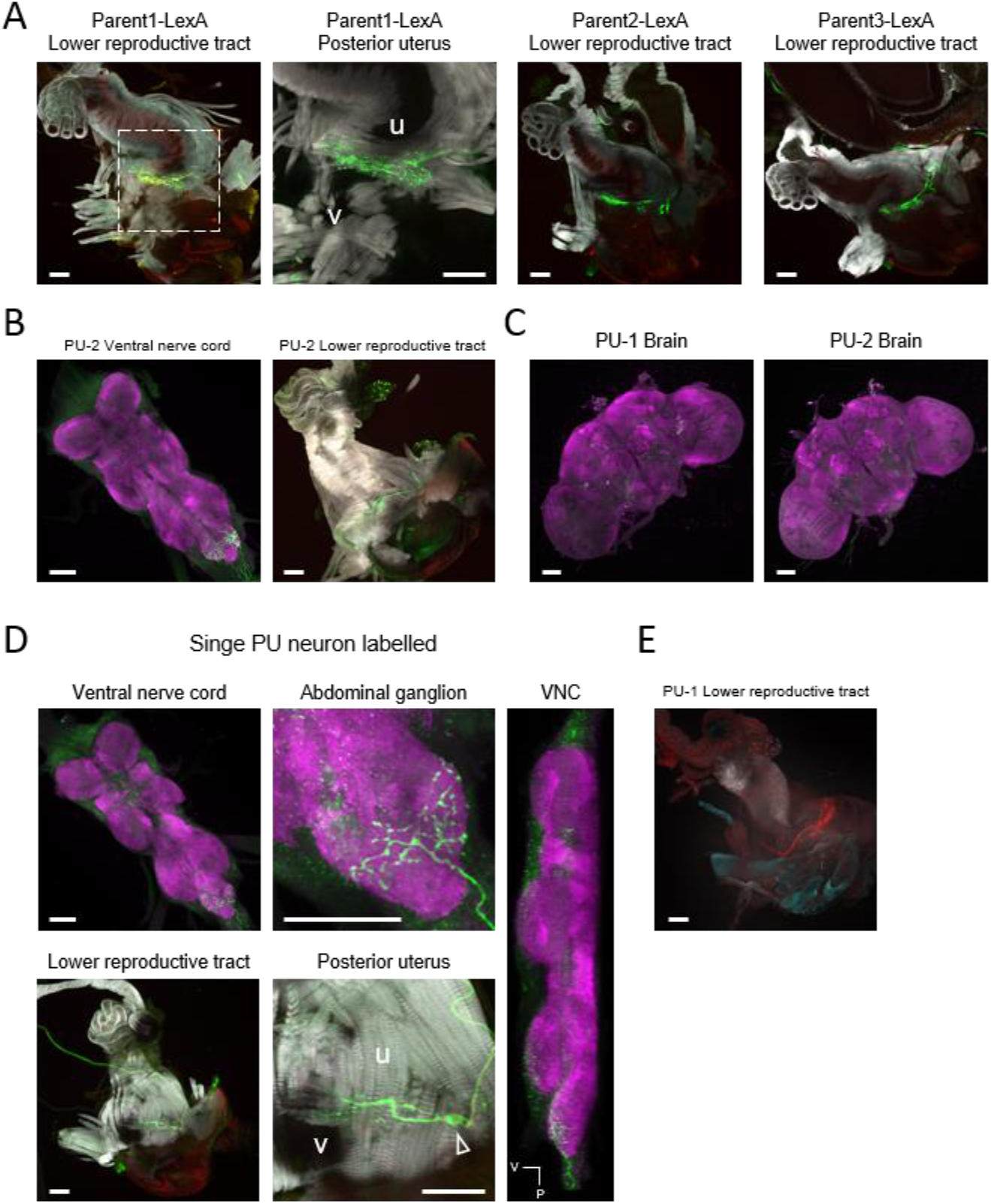
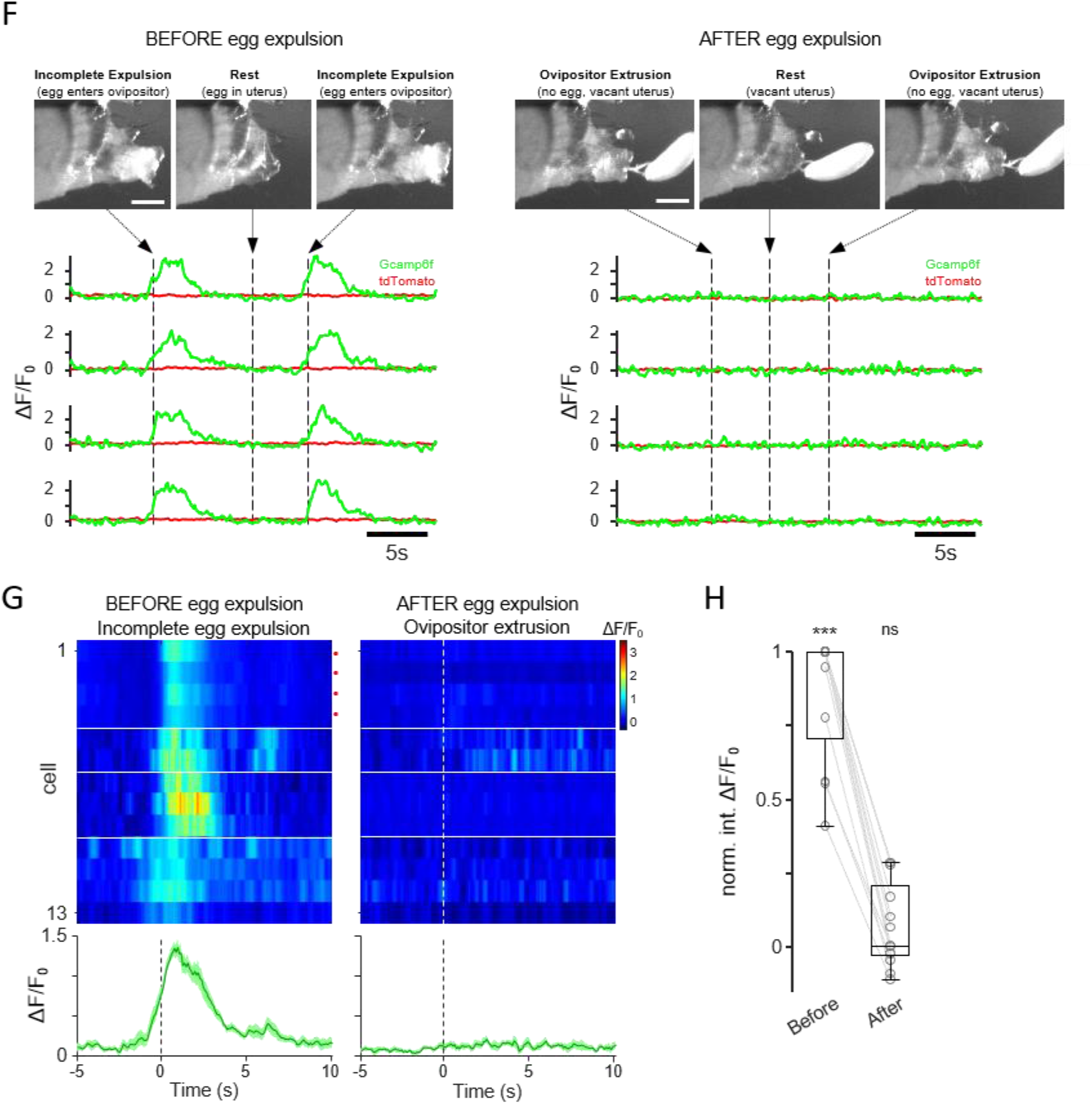
(A) Confocal images of the lower reproductive tract and posterior abdomen in three LexA lines that drive expression in PU neurons, stained with anti-GFP to reveal the membrane of labelled neurons (green) and anti-F-actin to visualize muscle fibers (gray). The abdominal cuticle is visualized by autofluorescence (red). Area in dashed box in the first image is enlarged in the second image; ‘u’, uterus; ‘v’, vagina. Scale bar is 50 μm. (B) Confocal images from a PU-2 > CD8-GFP female. Left, ventral nerve cord stained with anti-GFP (green) and nc82 (magenta). Right, lower reproductive tract and posterior abdomen stained with anti-GFP (green) and anti-F-actin (gray), and the abdominal cuticle visualized by autofluorescence (red). Scale bar is 50 μm. (C) Confocal images of the brain from a PU-1 > CD8-GFP female (left) and a PU-2 > CD8-GFP female (right), stained with anti-GFP (green) and nc82 (magenta). Scale bar is 50 μm. (D) Confocal images from a hs-FLP; PU-2 > (FRT.stop)CsChrimson female where stochastic labelling resulted in only a single PU neuron being labelled. Top two images and right image, ventral nerve cord stained with anti-GFP (green) and nc82 (magenta). Bottom two images, lower reproductive tract and posterior abdomen stained with anti-GFP (green) and anti-F-actin (gray), and the abdominal cuticle visualized by autofluorescence (red). The PU cell body is indicated in the second bottom image (“Posterior uterus”) by a white triangle; ‘u’, uterus; ‘v’, vagina. The right image (“VNC”) depicts a lateral projection of the ventral nerve cord after registration with a template; ‘V’, ventral; ‘P’, posterior. Scale bar is 50 μm. Flies with stochastic expression of CsChrimson were generated as described previously ^62^. (E) Confocal image of the lower reproductive tract and posterior abdomen of a PU-1 > DenMark female, stained with anti-DsRed to reveal the dendrites of labelled neurons (red), anti-F-actin (gray), and the abdominal cuticle visualized by autofluorescence (cyan). Scale bar is 50 μm. (F) Top left, video snapshots of the posterior abdomen depicting two incomplete egg expulsion events. Top right, snapshots depicting two ovipositor extrusion events after egg expulsion, and so lacking an egg. Scale bar is 200 μm. Bottom left and bottom right, two-photon calcium imaging of four PU axons, encompassing the above events, depicting relative fluorescence changes (ΔF/F0). Green traces, GCamp6f fluorescence; red traces, tdTomato anatomical marker. Arrows and vertical dashed lines indicate the corresponding time point for each video snapshot presented above. Note that the video snapshots were flipped in the vertical dimension here. (G) Relative fluorescence changes in PU neurons in response to incomplete expulsion events (left) and ovipositor extrusion events after egg expulsion (right), with individual responses shown above and the average response across all neurons shown below. Bottom, darker traces indicate mean response; lighter area represents SEM. The cells from (E) are indicated by a red dot. If multiple events occurred, the average response per neuron is presented here (n = 17, 1, 1, 5 incomplete expulsion events; n = 64, 1, 17, 2 ovipositor extrusion events after egg expulsion). Horizontal white lines demarcate recordings performed from different flies. Time = 0 is the time of event onset (see Methods). (H) Normalized population data, showing the 3-second integrated ΔF/F0 fluorescence levels during incomplete expulsion events (before egg expulsion) and ovipositor extrusion events (after egg expulsion). Statistical comparisons were made with the baseline, “0” value. ***p<0.001, ns p>0.05, Wilcoxon signed rank test.

**Figure S3.**
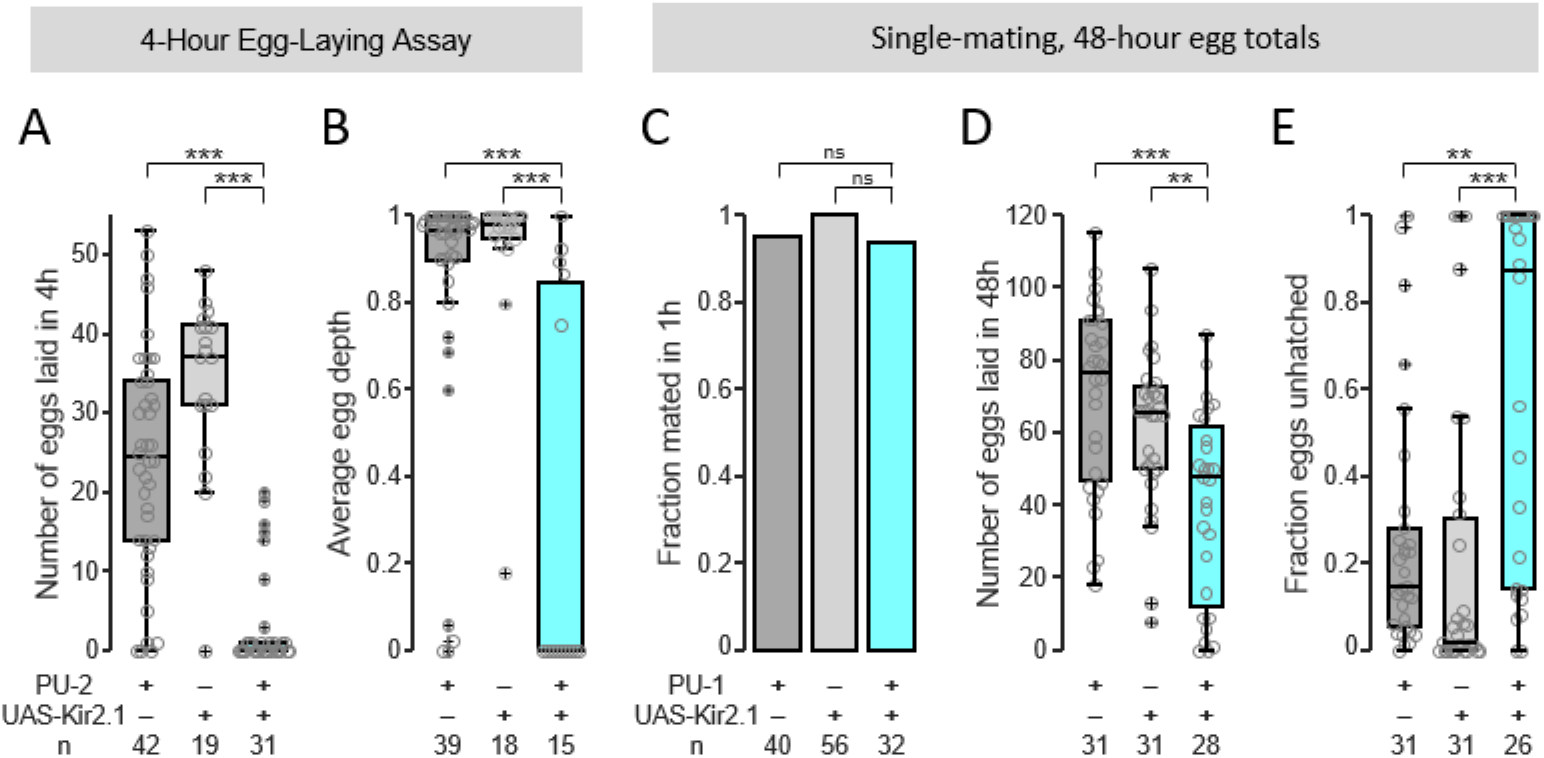
(A) Number of eggs laid on 1 % agar in a 4-hour window. For all panels in this figure, GAL4-only control is PU-1 > myr-GFP and UAS-only control is empty-SplitGAL4 > Kir2.1. For all panels in this figure except (C), *p<.05, **p<0.01, ***p<0.001, two-sided Wilcoxon rank sum test followed by Bonferroni correction. (B) Average depth of penetration of laid eggs (see Methods). (C) Fraction of virgin females that copulated within one hour. n.s, p>0.05, Fisher’s exact test. (D) Number of eggs laid in a 48-hour window following a single mating event. (E) Average fraction of eggs that did not hatch; from eggs counted in (D).

**Figure S4.**
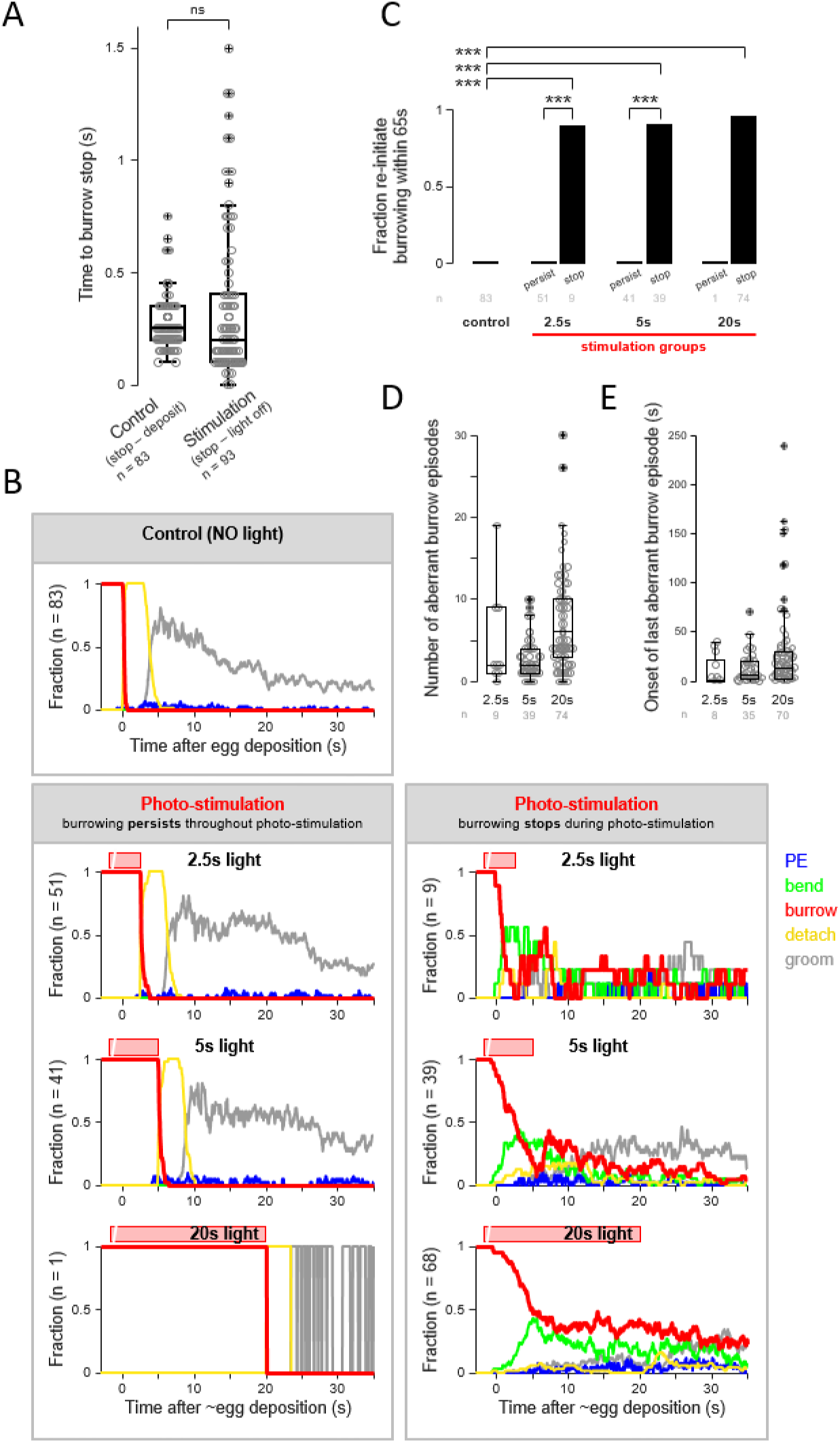
(A) Comparison of the timing that burrowing stopped after completed egg deposition in control flies with the timing that burrowing stopped after light offset in the subset of stimulation events where burrowing persisted throughout photo-stimulation. ns, p>0.05, two-sided Wilcoxon rank sum test. (B) Time course of the five annotated behaviors for the “no light” control (top) and three stimulus conditions (2.5s, 5s, and 20s light). For stimulation events, data are plotted separately for the events in which burrowing persisted throughout photo-stimulation (lower left), and the events in which burrowing stopped during photo-stimulation (lower right). The period of photo-stimulation is indicated by the red bar above each plot. (C) Fraction of events where burrowing was re-initiated within 65s of egg deposition for the “no light” control and three stimulus conditions (2.5s, 5s, and 20s light). For stimulation events, the fraction is plotted separately for the events in which burrowing persisted throughout photo-stimulation (“persist”), and the events in which burrowing stopped during photo-stimulation (“stop”). Events where burrowing was re-initiated within 65s of egg deposition were considered to have reverted to exploration instead of transitioning to the “reset” phase. ***p<0.001, Fisher’s exact test. (D) Number of aberrant burrowing episodes after egg deposition for the subset of events where burrowing stopped during photo-stimulation, separately plotted for all three stimulus conditions. An episode was considered aberrant if it occurred within 65 seconds of egg deposition or a previous aberrant episode (see Methods and Figure S4C). (E) The onset timing of the last aberrant burrowing episode expressed after egg deposition for the events where burrowing stopped during photo-stimulation and the fly reverted to exploration, separately plotted for all three stimulus conditions. The last aberrant episode after the fly reverted to exploration was determined as the first episode to precede a 65 second window devoid of burrowing behavior (see Methods).

